# Intravenous Administration of Serotonergic Psychedelics Produce Short-lasting Changes in Sleep-Wake Behavior and High Gamma Functional Connectivity in Rats

**DOI:** 10.1101/2025.10.12.681880

**Authors:** Nicholas Kolbman, Emma R. Huels, Amanda Nelson, Rachel Summerfield, Kanakaharini Byraju, Trent Groenhout, Tiecheng Liu, Anthony G. Hudetz, Giancarlo Vanini, Dinesh Pal

## Abstract

**Background and Purpose:** Given the increase in recreational psychedelic use and ongoing efforts to explore psychedelics as therapeutic agents for mental health disorders, there is an urgent need to understand the effect of psychedelics such as psilocybin and N,N-dimethyltryptamine (DMT) on sleep-wake states, which share a bidirectional relationship with mental health. Here, we investigated the effects of intravenous psilocybin and DMT on sleep-wake states and EEG spectral power and functional connectivity in rats.

**Experimental Approach:** Sprague Dawley rats (n=25, 13 male) were surgically instrumented to record high-density EEG (27 electrodes) and EMG during 12-h light and 12-h dark cycle after intravenous psilocybin (2.5 mg/kg, 10 mg/kg), DMT (3.75 mg/kg, 7.5 mg/kg) or 0.9% saline. The EEG/EMG data were scored in 4-second epochs into wake, slow-wave sleep (SWS), and rapid eye movement (REM) sleep. EEG spectral power and corticocortical coherence, a surrogate for functional connectivity, were computed in 12-second epochs.

**Key Results:** Psilocybin and DMT delayed the onset of SWS and REM sleep, and caused a short-lasting increase in wakefulness and decrease in SWS. Psilocybin also produced a 1) decrease in REM sleep, 2) decrease in theta power and coherence and increase in high gamma power and coherence during wake and SWS, and 3) increase in high gamma coherence during REM sleep. DMT increased gamma coherence only during wakefulness.

**Conclusions and Implications:** Serotonergic psychedelics have minimal effects on sleep-wake states. The enhanced high gamma functional connectivity suggests that the psychedelic-induced changes in EEG/neural dynamics can occur independent of the arousal states.

## Introduction

There is a growing interest in serotonergic psychedelics, including psilocybin and N,N-dimethyltryptamine (DMT), as potential therapeutic agents for neuropsychiatric illnesses such as anxiety, depression, and substance use disorders.^1–5^ There has also been concerted efforts towards decriminalization of serotonergic psychedelics for recreational use.^6^ With the use of serotonergic psychedelics becoming more prevalent, there is a critical need to understand their effect on sleep-wake behavior, which is intricately linked to general well-being and shares a bidirectional relationship with neuropsychiatric illnesses.^7–9^

While anecdotal reports suggest sleep disturbances following consumption of serotonergic psychedelics, the results from survey studies remain mixed, with some reporting sleep disturbances and others improved sleep.^10–12^ More recently, peroral psilocybin administration in human subjects was shown to increase the latency to the onset of nightly rapid eye movement (REM) sleep without any statistically significant alteration in the time spent in different sleep-wake states.^13^ Furthermore, few studies have systematically assessed the effect of serotonergic psychedelics on sleep-wake states in rodent models, with several dating back fifty years and limited by their methodological approaches, such as the use of sparse electroencephalogram (EEG) recordings that were limited to low frequency oscillations (<40 Hz), and routes of administration lacking complete drug bioavalability.^14–17^ In a recent study, intraperitoneal administration of psilocin — a direct metabolite and the psychoactive product of psilocybin — in mice produced an increase in the latency to the onset of REM sleep and a transient increase in wakefulness and decrease in slow-wave sleep (SWS).^18^ Similar effects — i.e., increased wakefulness, disrupted SWS, and delayed onset of REM sleep — have also been reported after intraperitoneal administration of the synthetic psychedelic and 5-HT2A receptor agonist, 2,5-dimethoxy-4-iodoampethamine (DOI), in rats.^19^

Despite these previous findings, two key gaps in our understanding of the effect of serotonergic psychedelics on sleep-wake states remain. *First,* the effects of psilocybin and DMT on sleep-wake behavior and cortical neurophysiological measures across sleep-wake states in a rat model are unknown. *Second*, the previous rodent studies have employed either intraperitoneal or subcutaneous routes of administration, which, as opposed to an intravenous route, have incomplete bioavailability and thus have limited translational applicability. Therefore, in this study, we investigated the dose-dependent effects of intravenously administered psilocybin and DMT on sleep-wake behavior and measures of spectral power and functional connectivity during sleep-wake states in adult male and female rats.

## Methods

### Subjects

All experimental procedures were approved by the Institutional Animal Care and Use Committee at the University of Michigan and were conducted in compliance with the Guide for the Care and Use of Laboratory Animals (National Academies Press, 8th Edition, Washington DC, 2011). The experiments were conducted in adult Sprague Dawley rats (300-350 g, Charles River Laboratories Inc., MA) of both sexes (n=25, 13 male). The rats were housed in a temperature-controlled facility, provided with ad libitum food and water, and maintained on a 12 h:12 h light-dark cycle (lights on at: 08:00 am).

### Surgical procedures

The rats were placed in an air-tight clear rectangular chamber (10.0 inches×4.8 inches×4.2 inches) to induce general anesthesia with 4–5% isoflurane (Piramal Enterprises, Telangana, India) in 100% oxygen. After anesthetic induction, the cranial surface between the eyes and neck was disinfected and shaved, and the rats were positioned in a stereotaxic frame (Model 963, David Kopf Instruments, Tujunga, CA) using blunt ear bars. Isoflurane during surgery was delivered via a rat nose cone (Model 906, David Kopf Instruments, Tujunga, CA) mounted on a stereotaxic frame and was titrated (1–2%) to maintain absence of pedal withdrawal reflex. The anesthetic concentration was continuously monitored using an anesthetic agent analyzer (Datex Medical Instrumentation, Tewksbury, MA). Body temperature was monitored using a rectal probe (RET-2 ISO, Physitemp Instruments, Inc., Clifton, NJ) and maintained at 37.0 ± 1 ◦C using a small animal far-infrared heating pad (Kent Scientific Co., Torrington, Connecticut). The rats were divided into two groups and were implanted with an indwelling chronic catheter (Micro-Renathane tubing, MRE-040, Braintree Scientific, MA) in the jugular vein. One group of rats received either 0.9% saline (vehicle control) or two different doses of psilocybin (Cayman Chemical, product no. 14041) whereas the other group received either 0.9% saline or two different doses of N,N-Dimethyltryptamine fumarate (DMT) (Cayman Chemical, product no. 9003568). The catheter was tunneled under the skin, mated with a pin-port (PNP3M-F22, Instech Laboratories Inc., Plymouth Meeting, PA), and affixed on top of the skull. Under aseptic conditions, the cranium was exposed, and 27 stainless-steel screw electrodes were implanted across frontal, parietal, and occipital cortices to acquire high-density EEG data. A stainless-steel electrode was implanted over the nasal sinus to serve as a reference electrode, and a pair of insulated (except at the tips) wires (Cat # 791400, A-M Systems, Sequim, WA) were inserted into the dorsal nuchal muscles to record electromyogram (EMG) activity. The free ends of the EEG and EMG electrodes were soldered into a 32-pin connector (P/N: 853-43-100-10-001000, Mill-Max, Mansfield, TX) and the entire assembly was affixed to the cranial surface using dental cement (51459, Stoelting Dental Cement, Wood Dale, IL). All rats received carprofen (5 mg/kg, s.c.) and buprenorphine (0.01 mg/kg, s.c., Buprenex, Reckitt Benckiser Pharmaceuticals, Richmond, VA) for pre-surgical analgesia, and cefazolin (West-Ward-Pharmaceutical, Eatontown, NJ) (20 mg/kg, s.c.) as pre-surgical antibiotic. The rats also received buprenorphine (0.03 mg/kg, s.c.) every 8–12 h for 48 h for post-surgical analgesia. To maintain catheter patency, the jugular venous catheter was flushed with 0.2 mL of heparinized (1 unit/mL, Sagent Pharmaceuticals, Schaumburg, IL) saline and locked with 0.05 mL of Taurolidine-Citrate Catheter lock solution (TCS-04, Access Technologies, Skokie, IL) every 5-7 days. The rats were provided 2-3 weeks of post-surgical recovery.

### Experimental design

The experimental design is illustrated in **figure 1**. After a week of post-surgical recovery, the rats were continuously conditioned for at least 1-2 weeks to the sleep-wake recording set-up, during which the rats were handled daily and connected to the EEG/EMG recording cables in the recording chambers. The rats were connected to the EEG/EMG recording system at least 12-16 h before the start of EEG/EMG recording at the beginning of lights-ON period. EEG and EMG signals were recorded using a Cereplex Direct recording system (Blackrock Microsystems, Salt Lake City, UT). Monopolar EEG (0.1-300 Hz, sampling rate 1 kHz) was recorded from all electrodes except three, which were arranged in a bipolar montage (frontal-frontal, frontal-parietal, and parietal-parietal, 0.1-125 Hz, sampling rate 500 Hz) for polysomnography. The EMG was bandpass filtered between 0.1 Hz and 125 Hz and sampled at 500 Hz. The recording headstage was equipped with a motion sensor from which the data were bandpass filtered between 0.1 Hz and 50 Hz and sampled at 500 Hz. The signals from the motion sensor were used as an additional aid for identification of wake epochs. After about an hour of EEG/EMG recording in the lights-ON period, the rats (n=6 male, 6 female) received a single intravenous bolus of either 0.9% saline (vehicle control) or one of the two doses of psilocybin (low dose: 2.5 mg/kg, high dose: 10 mg/kg), with the exception of one rat that did not undergo the saline experiment due to health concerns. Another group of rats (n=7 male, 6 female), prepared similarly for polysomnography, received either 0.9% saline (vehicle control) or one of the two doses of DMT (low dose: 3.75 mg/kg, high dose: 7.5 mg/kg). The high-density EEG and EMG data collection continued uninterrupted for the next 24 hours.

**Figure 1.**
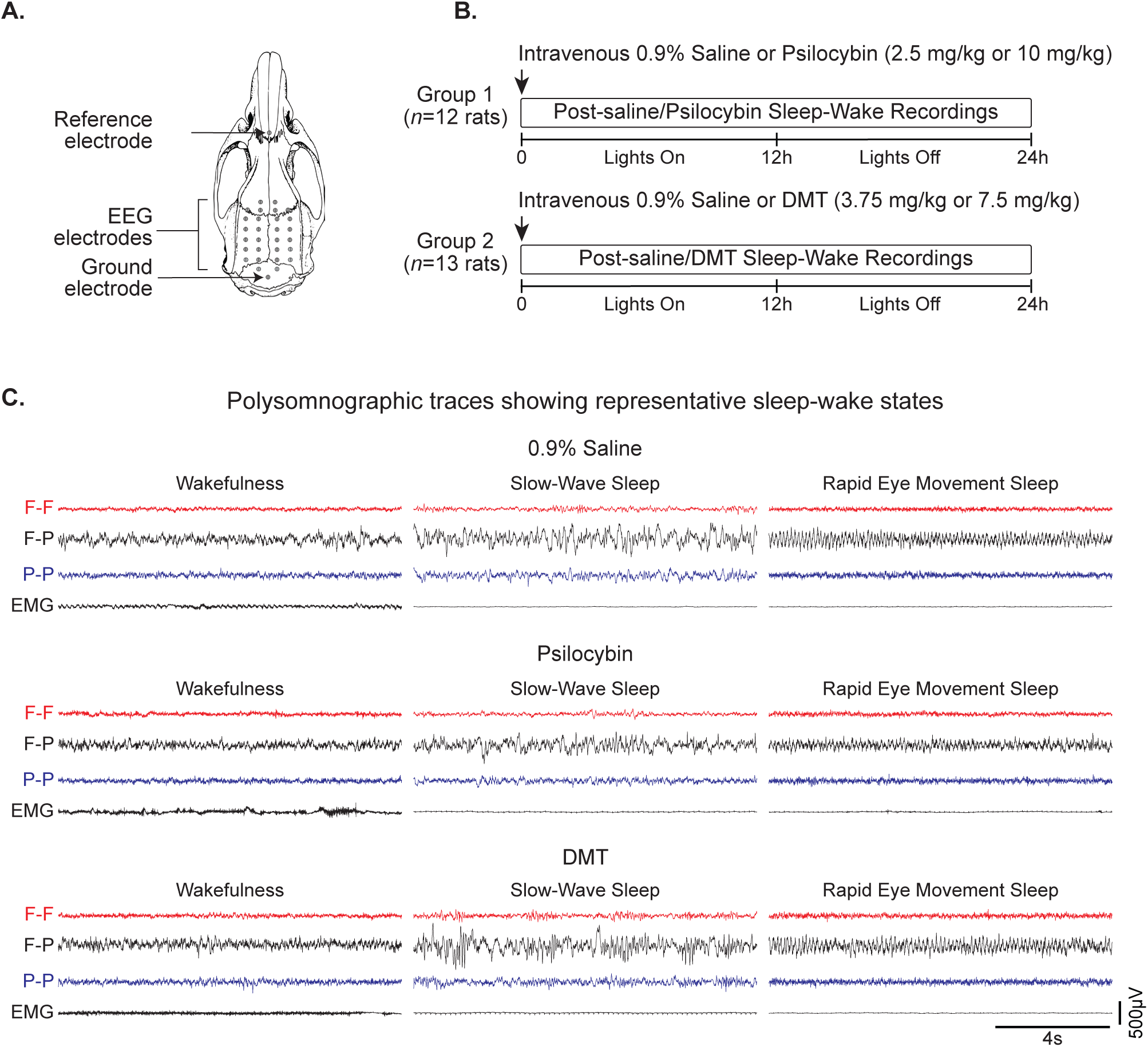
Schematic showing experimental design and timeline. **A.** Sprague Dawley rats were surgically instrumented with stainless steel screw electrodes for high-density EEG recording (27 channels), bilateral wire electrodes in the nuchal muscle to record electromyogram, and an indwelling chronic catheter in the jugular vein. **B.** Schematic showing experiment timeline and treatment groups. One subgroup of the rats (n=12, 6 male) received 0.9% saline, 2.5 mg/kg psilocybin, and 10 mg/kg psilocybin through the jugular vein catheter on different days separated by at least 5-7 days and in a randomized order. In a similar manner, the other subgroup (n=13, 7 male) received 0.9% saline, 3.75 mg/kg DMT, and 7.5 mg/kg N,N-dimethyltryptamine (DMT). All infusions were done at the beginning of the light cycle. **C.** EEG and EMG traces show representative sleep-wake states. F – frontal EEG, P – Parietal EEG,

### Sleep-wake state identification

EEG and EMG data were scored manually using SleepSign (Kissei Comtec Inc.) in 4-second epochs into the following sleep-wake states: wake state characterized by low-amplitude fast EEG along with high muscle tone, slow-wave sleep (SWS) characterized by high-amplitude slow EEG along with low muscle tone, and rapid eye movement (REM) sleep characterized by low-amplitude fast EEG along with muscle atonia. The percentage of time spent in each state, the number of epochs per state, and the mean duration per episode for each state after 0.9% saline, psilocybin, or DMT infusion were calculated and averaged into 2 h bins for the entire post-infusion period. Latency to the onset of SWS and REM sleep were quantified as the occurrence of the first SWS and REM sleep episode following the infusion of 0.9% saline, psilocybin, or DMT.

### EEG analysis

The EEG signals from all except one male and one female rat in the psilocybin group were used for the power spectral density and coherence analyses. All analyses were conducted using MATLAB. The analyses were focused on the first four hours of EEG data after the start of the drug/saline infusion and were performed on three consecutive 4-second artifact-free non-overlapping windows (totaling a 12-second window) for each of the sleep-wake states (wake, SWS, REM sleep). Power spectral density (PSD) in 0.5-200 Hz range was estimated for each channel using MATLAB’s implementation of Welch’s method (pwelch.m) (4-second windows, 2-second overlap). Relative power was then computed for each frequency for each 12-second window by dividing a given frequency’s absolute power by the total absolute power across all frequencies (0.5-200 Hz). Functional connectivity was calculated by computing the magnitude-squared coherence between each electrode pair (e.g., electrode 1 vs. electrode 2, electrode 1 vs. electrode 3, and so for all pair of electrodes in the entire montage) using ‘mscohere.m’ in the same frequency range (0.5-200 Hz) and the same analysis window (12-seconds) as were used for the power spectral analysis. PSD and coherence values were then averaged across the following frequency bands: delta (1-4 Hz), theta (4-10 Hz), medium gamma (70-110 Hz), and high gamma (125-155 Hz) bands. These frequency bands were selected based on previous studies from our and other laboratories that have shown their involvement in states of consciousness.^20–22^ PSD and coherence values were averaged across time (1 hour bins) and electrodes for the mean global relative power and mean global coherence.

### Statistical analysis

All statistical analyses were conducted using R software version 4.3.1 (R Core Team, 2023) and in consultation with the *Consulting for Statistics, Computing, and Analytics Research* core at the University of Michigan. GraphPad Prism software version 9.5 (GraphPad Software, San Diego, CA, United States) was used to create all graphs. Statistical comparisons were conducted in 2 h bins for sleep-wake behavior. A linear mixed model was fit with lme4 to compare the effects of administration of drugs (psilocybin or DMT) with vehicle control (0.9% saline) on the 1) percentage of time spent in sleep-wake states, 2) mean number of epochs for each state, and 3) mean epoch duration for each state. The linear mixed model included dose (0.9% saline, 2.5 mg/kg psilocybin, and 10 mg/kg psilocybin; or 0.9% saline, 3.75 mg/kg DMT, and 7.5 mg/kg DMT), time (post-infusion 2-24 h), and their interaction as fixed effects, with the subject (rat) serving as a random intercept. A similar model was used to compare drug and vehicle effects on the latency to the onset of SWS and REM sleep, where dose was again included as a fixed effect and the subject (rat) served as a random intercept. Statistical comparisons for EEG analyses were conducted in 1 h bins. A linear mixed model was fit to compare drug effects (psilocybin or DMT) with vehicle control (0.9% saline) on spectral power and functional connectivity within the delta, theta, medium gamma, and high gamma bands for each sleep-wake state (wake, SWS, REM sleep). The linear mixed model included dose, time (post-infusion 1-4 h), and their interaction as fixed effects, with the subject (rat) serving as a random intercept. All data are reported as mean ± standard error of the mean (s.e.m.). Post-hoc comparisons between saline and drug conditions were extracted, the alpha threshold was set to p < 0.05, and associated p-values were adjusted with Tukey’s method.

## Results

### Effect of Intravenous Administration of Psilocybin or DMT on Sleep-Wake Behavior

As compared to the saline infusion, intravenous infusion of psilocybin produced a dose-dependent increase in the mean latency to the onset of REM sleep (low dose: p=0.03, high dose: p<0.0001) (**Figure 2A**). Psilocybin also produced a statistically significant increase in the mean latency to onset of SWS, but only at the low dose (p=0.002) (**Figure 2B**). Similar effects were observed after DMT administration, such that infusion of high-dose DMT increased the mean latency to the onset of REM sleep (p=0.0003) (**Figure 2C**). Additionally, infusion of low-dose (p=0.003) or high-dose (p<0.0001) DMT increased the mean latency to the onset of SWS (**Figure 2D**).

**Figure 2.**
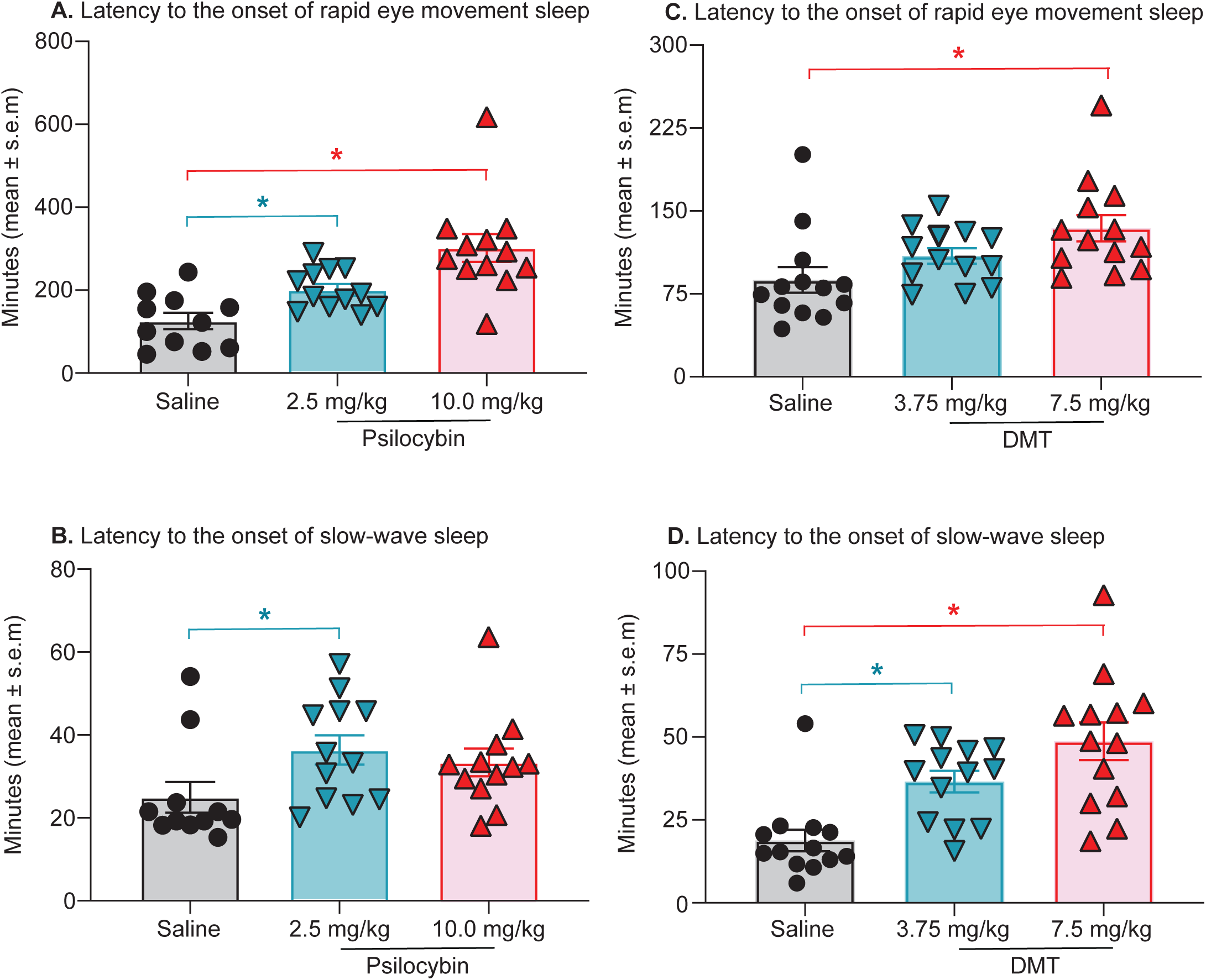
Intravenous administration of psilocybin or N,N-dimethyltryptamine (DMT) increased the latency to the onset of rapid eye movement sleep and slow-wave sleep. **A.** As compared to intravenous saline infusion, intravenous infusion of 2.5 mg/kg psilocybin (p=0.03) or 10 mg/kg psilocybin (p<0.0001) significantly increased the latency to the onset of rapid eye movement sleep. **B.** As compared to saline infusion, delivery of 2.5 mg/kg psilocybin (p=0.002), but not 10 mg/kg psilocybin (p=0.09), significantly increased the latency to the onset slow-wave sleep. **C.** As compared to intravenous saline infusion, intravenous infusion of 7.5 mg/kg DMT (p=0.0003), but not 3.75 mg/kg DMT (p=0.09) significantly increased the latency to the onset of rapid eye movement sleep. **D.** As compared to intravenous saline infusion, intravenous infusion of 3.75 mg/kg DMT (p=0.003) or 7.5 mg/kg DMT (p<0.0001) significantly increased the latency to the onset of slow-wave sleep. The significance symbols denote p<0.05 and are color coded to match the symbols/bars for each dose. *Significant as compared to saline infusion.

Intravenous administration of low-dose (p=0.0002) or high-dose (p=0.006) psilocybin produced a statistically significant increase in the time spent in wake state during the first 2h in the post-infusion period (i.e., hours 0-2 during the light phase) (**Figure 3A**). As compared to saline infusion, infusion of low-dose psilocybin produced a decrease in the time spent awake in post-infusion hours 16-18 (p=0.002) and hours 22-24 (p=0.02); a similar decrease was observed in post-infusion hours 16-18 (p=0.0005) and 18-20 (p=0.02) after high-dose psilocybin administration (**Figure 3A**). Concurrently, a decrease in SWS during the first 2h bin in the post-infusion period was observed after the administration of low-dose (p<0.0001) or high-dose (p=0.002) psilocybin (**Figure 3B**). There was also a significant increase in the time spent in SWS during the post-infusion hours 16-18 for both the low-dose (=0.007) and the high-dose (p=0.002) psilocybin groups (**Figure 3B**) and during post-infusion hours 22-24 (p=0.03) after low-dose psilocybin infusion (**Figure 3B**). As compared to saline infusion, infusion of high-dose psilocybin decreased the time spent in REM sleep during post-infusion hours 2-4 (p=0.04) and hours 4-6 (p=0.03) while both low-dose and high-dose psilocybin increased the time spent in REM sleep during post-infusion hours 16-18 (low dose: p=0.01; high dose: p=0.01) (**Figure 3C**).

**Figure 3.**
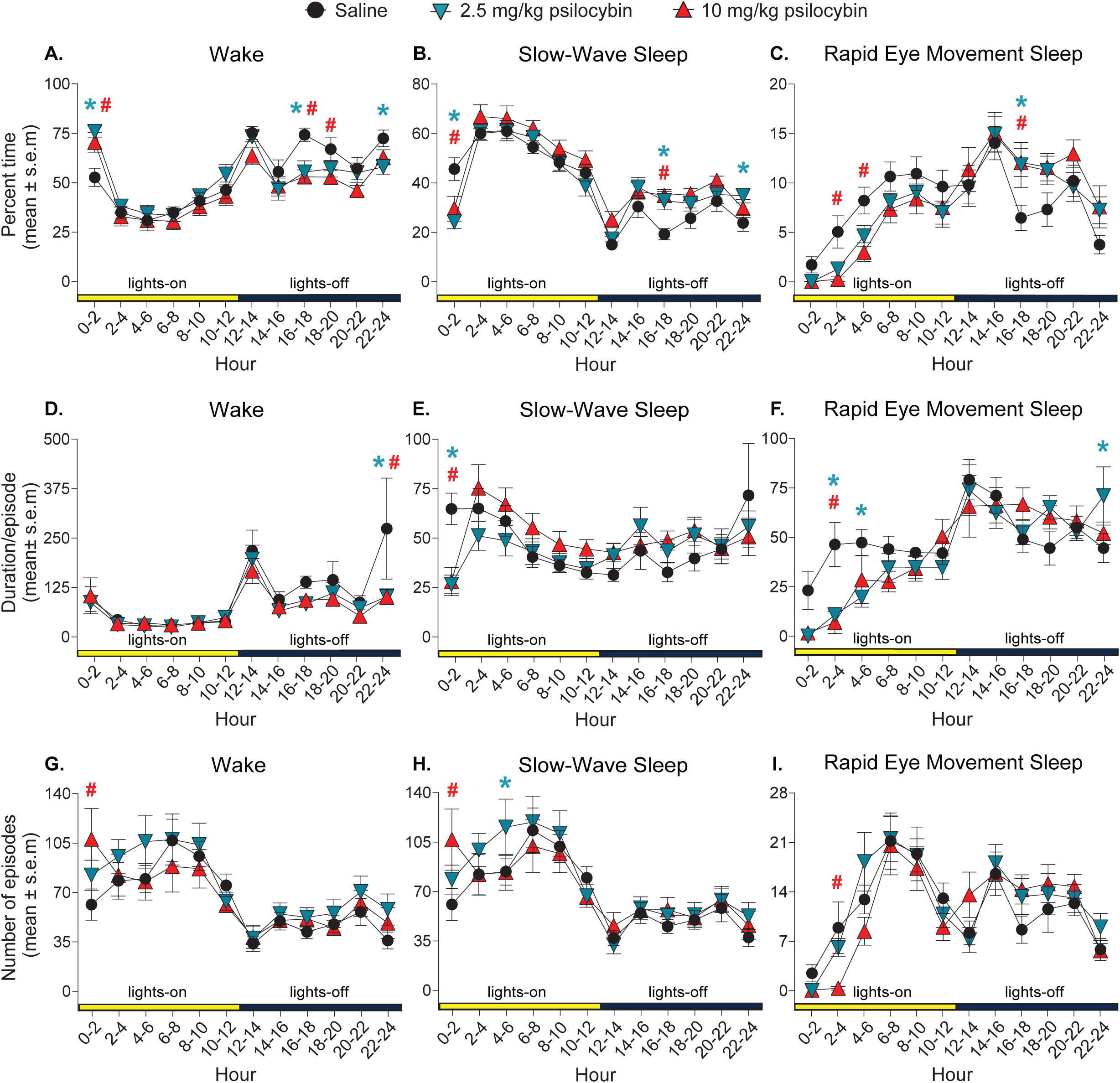
Intravenous administration of psilocybin produces a short-lasting increase in wakefulness, decrease in slow-wave sleep, and suppression of rapid eye movement sleep. The percent time spent in wake state (**A**.), slow-wave sleep (**B.**), and rapid eye movement sleep (**C.**) after intravenous infusion of 0.9% saline (black circles), 2.5 mg/kg psilocybin (teal inverted triangles), and 10 mg/kg psilocybin (red triangles). Panels **D.-F.** show the effect of saline or psilocybin infusion on the mean duration per episode during wake, slow-wave sleep, and rapid eye movement sleep. The effect of saline or psilocybin infusion on the mean number of episodes across sleep-wake states is shown in panels **G.-I.** Asterisk and pound signs indicate statistical significance (p<0.05) for the comparison between low dose (2.5 mg/kg) and saline infusion, and high dose (10 mg/kg) and saline infusion, respectively.

Although there was no change in the mean duration of wake epochs in the lights-on period immediately after low-dose or high-dose psilocybin administration (**Figure 3D**), there was a significant increase in the mean number of wake episodes (0h-2h: p=0.0009) (**Figure 3G**) during the first 2h block in the post-infusion period, which accounts for the increase in time spent in wakefulness after the infusion of high-dose psilocybin. The decrease in time spent in SWS during the first 2h in the post-infusion period after low-dose (p=0.0002) or high-dose (p=0.0004) psilocybin was due to a significant decrease in the mean duration of SWS episodes (**Figure 3E**). Additionally, there was a statistically significant increase in the mean number of SWS episodes during the 0h-2h bin (compared to saline, p=0.0008) after the infusion of high-dose psilocybin, and in the 4h-6h bin after low-dose psilocybin infusion (p=0.03) (**Figure 3H**). The decrease in REM sleep after administration of high-dose psilocybin appears to be driven by the concurrent decrease in the mean duration of REM sleep episodes (p=0.0005) as well as a decrease in the mean number of REM sleep epochs (p=0.02) in the 2-4h time bin (**Figure 3F & I**). Although the mean time spent in REM sleep was not statistically significantly different between low-dose psilocybin and saline group at 0h-6h post-infusion, there appears to be a trend towards an overall decrease in the time spent in REM sleep between hours two and six (**Figure 3C**). This trend appears to be driven by a reduction in the mean duration of REM sleep episodes for the low-dose psilocybin group (2h-4h: p=0.002; 4h-6h: p=0.02) (**Figure 3F**).

As compared to saline infusion, infusion of high-dose DMT significantly increased the time spent in wake (p=0.007) and decreased the time spent in SWS (p=0.001) during the first 2h bin in the post-infusion period (**Figure 4A & B**). Following the onset of the dark phase, both low-dose and high-dose DMT decreased the time spent awake (low dose: p=0.01; high dose: p=0.004) and increased the time spent in SWS (low dose: p=0.03; high dose: p=0.04) and REM sleep (low dose: p=0.04; high dose: p=0.0004) in the 14-16h time bin post-DMT infusion (**Figure 4A-C**). The time spent in REM sleep remained increased in the 16-18h time bin for the low-dose DMT group (p=0.01) (**Figure 4C**).

**Figure 4.**
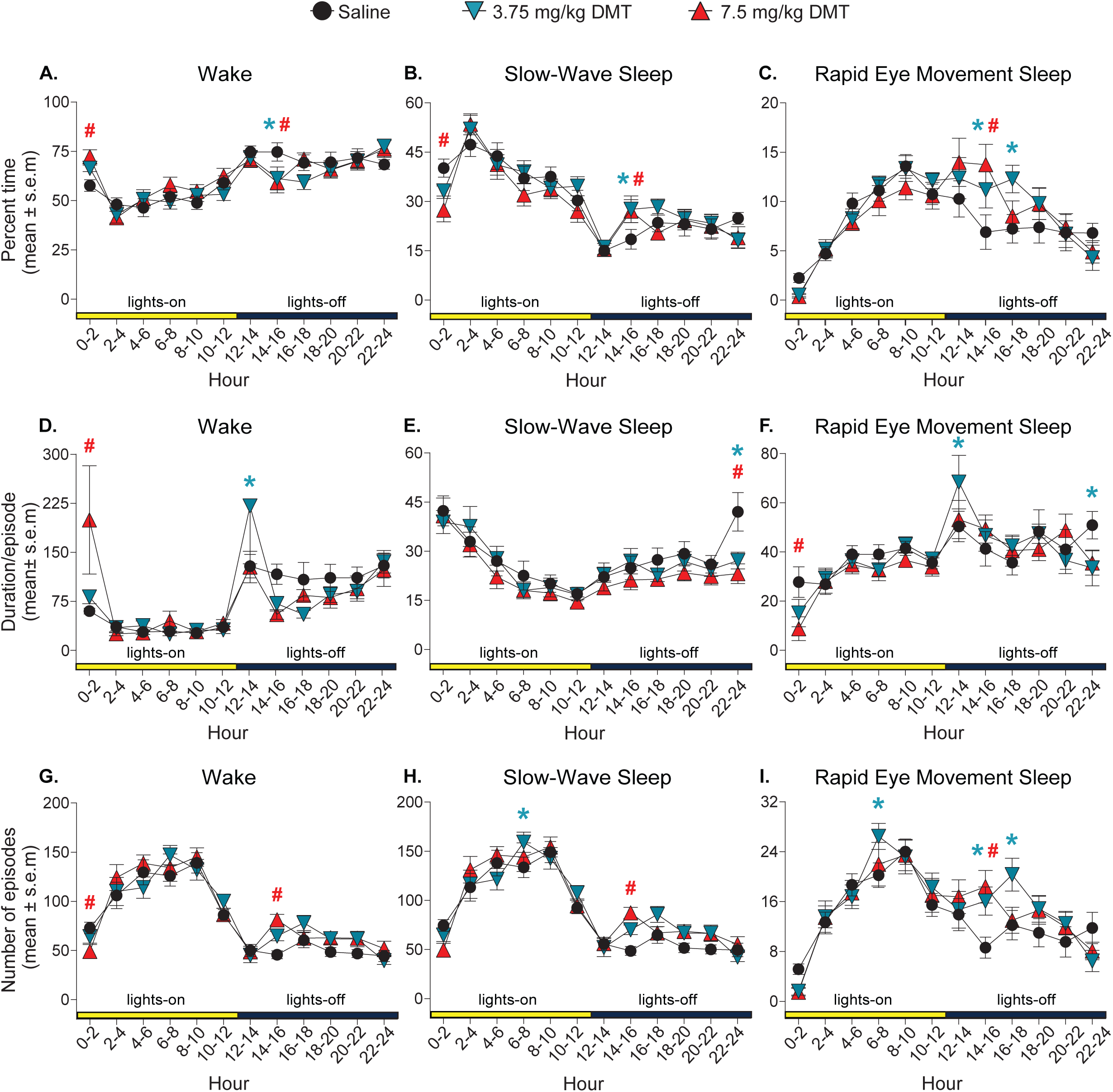
Intravenous administration of N,N-dimethyltryptamine (DMT) produces a short-lasting increase in wakefulness and decrease in slow-wave sleep. The percent time spent in wake state (**A**.), slow-wave sleep (**B.**), and rapid eye movement sleep (**C.**) after intravenous infusion of 0.9% saline (black circles), 3.75 mg/kg DMT (teal inverted triangles), and 7.5 mg/kg DMT (red triangles). Panels **D.-F.** show the effect of saline or DMT infusion on the mean duration per episode during wake, slow-wave sleep, and rapid eye movement sleep. The effect of saline or DMT infusion on the mean number of episodes across sleep-wake states is shown in panels **G.-I.** Asterisk and pound signs indicate statistical significance (p<0.05) for the comparison between low dose (3.75 mg/kg) and saline infusion, and high dose (7.5 mg/kg) and saline infusion, respectively.

The acute increase in wakefulness in the first 2h time bin after the infusion of high-dose DMT appears to be driven by a significant increase in the mean duration of wake episodes (p<0.0001) (**Figure 4D**. In contrast, there was no change in the duration of SWS episodes in the first 2h time bin following high-dose DMT (p>0.05) (**Figure 4E**) and a significant decrease in the duration of REM sleep episodes in the same time bin (p=0.02) (**Figure 4F**). Following the onset of the dark phase, there was an increase in the mean duration of wake (p=0.01) and REM sleep (p=0.03) episodes in the 12h-14h time bin in the post-infusion period following low-dose DMT (**Figure 4D & F**). The mean duration of REM sleep episodes subsequently decreased in the 22-24h time bin following low-dose DMT (p=0.03) (**Figure 4F**), as did the mean duration of SWS episodes following either dose of DMT for the same time bin (low dose: p<0.0001; high dose: p<0.0001) (**Figure 4E**). During the light phase, there was an increase in the number of SWS (p=0.04) and REM sleep (p=0.04) epochs in the 6h-8h post-infusion time bin for the low-dose group (**Figure 4H & I**). Following the onset of the dark phase, there was an increase in the number of wake (p=0.002) and SWS (p=0.001) epochs following high-dose DMT in the 14h-16h post-infusion time bin, as well as an increase in the number of REM sleep epochs following either dose in the same time bin (low dose: p=0.01; high dose: p=0.0008) (**Figure 4G-I**). The number of REM sleep epochs remained increased in the 16h-18h time bin following low-dose DMT (p=0.006) (**Figure 4I**).

### Effect of Intravenous Administration of Psilocybin or DMT on Spectral Power Across Sleep-Wake States

The effects of psilocybin or DMT infusion on the global relative spectral power were analyzed during the first four hours post-infusion in four frequency bands that have been shown to be consistently associated with states of arousal: delta (1-4 Hz), theta (4-10 Hz), medium gamma (70-110 Hz), and high gamma (125-155 Hz).^20^ The most salient effect of psilocybin on spectral power during wakefulness was observed in the medium gamma band, which showed a significant increase in power as compared to that observed after saline infusion. The increase in medium gamma power during wakefulness was observed after the administration of both low-dose (hour 1: p<0.0001, hour 2: p<0.0001, hour 3: p<0.0001, hour 4: p=0.003) and high-dose (hour 1: p<0.0001, hour 2: p=0.003, hour 3: p=0.005, hour 4: p<0.0001) psilocybin (**Figure 5A**). Delta power during wakefulness also increased in the first post-infusion hour after the administration of low-dose (p=0.02) and high-dose (p=0.0002) psilocybin (**Figure 5A**). Psilocybin administration at both the low dose (p=0.0002) and high dose (p<0.0001) decreased theta power during wake in the first post-infusion hour, whereas only high-dose psilocybin produced a similar decrease in post-infusion hour 2 (p=0.03) and hour 3 (p=0.005) (**Figure 5A**). High gamma power during wake state also increased in all four hours (hour 1: p=0.0003, hour 2: p=0.01, hour 3: p=0.009, hour 4: p=0.005) after administration of low-dose psilocybin and increased during hour 4 after administration of high-dose psilocybin(p=0.01) (**Figure 5A**).

**Figure 5.**
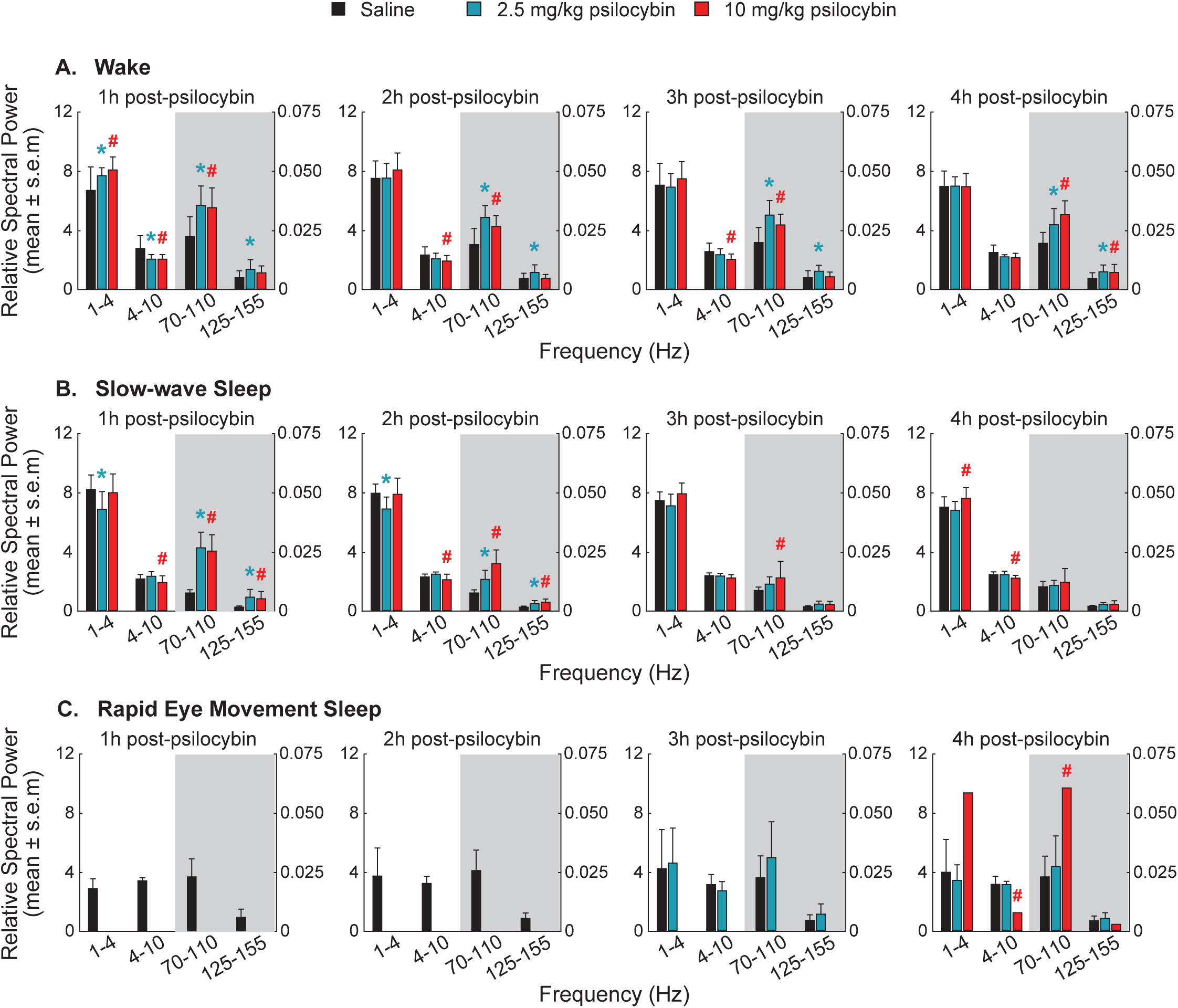
Effect of intravenous administration of psilocybin on power spectral density across sleep-wake states. Power spectral density analysis of the EEG from first four post-infusion (0.9% saline and psilocybin) hours. Relative power values were averaged across the delta (1-4 Hz), theta (4-10 Hz), medium gamma (70-110 Hz), and high gamma (125-155 Hz) frequency bands. The effect of intravenous administration of 0.9% saline (black bars), 2.5 mg/kg psilocybin (teal bars), and 10 mg/kg psilocybin (red bars) on the average global relative power during post-infusion hours 1 through 4 is shown for the wake state (**A.**), slow-wave sleep (**B.**), and rapid eye movement sleep (**C.**) For each plot, the left y-axis corresponds to values for the delta and theta bands whereas the right y-axis corresponds to values for the medium and high gamma bands. The significance symbols denote p<0.05. Asterisk and pound signs indicate statistical significance (p<0.05) for the comparison between low dose psilocybin (2.5 mg/kg) and saline infusion, and high dose psilocybin (10 mg/kg) and saline infusion, respectively.

After low-dose psilocybin infusion, delta power during SWS decreased during the first two hours after infusion (hour 1: p<0.0001, hour 2: p=0.002) and increased for the high-dose group during hour 4 (p=0.03) (**Figure 5B**). Theta power during SWS decreased in the high-dose psilocybin group during hours 1, 2, and 4 (hour 1: p=0.008, hour 2: p=0.04, hour 4: p=0.008) (**Figure 5B**) Medium gamma relative power during SWS increased in both the low-dose and high-dose psilocybin groups for hour 1 (low dose: p<0.0001, high dose: p<0.0001) and hour 2 (low dose: p=0.004, high dose: p<0.0001), whereas only high-dose psilocybin infusion showed an increase in hour 3 (p=0.005) (**Figure 5B**). During SWS, high gamma relative power increased for both the low-dose and high-dose groups for hour 1 (low dose: p<0.0001, high dose p<0.0001) and hour 2 (low dose: p=0.02, high dose: p=0.0007) (**Figure 5B**).

Due to the suppression of REM sleep after psilocybin infusion, the analyzable REM sleep epochs were found only in hour 4, during which theta relative power decreased (p=0.0009) and medium gamma relative power increased (p=0.009) after high-dose psilocybin; there were no statistical changes in the delta or high gamma bands during REM sleep following either of the psilocybin doses (**Figure 5C**).

Changes in relative power following intravenous DMT were limited. The only effects of intravenous DMT infusion on relative power during wake for any of the four frequency bands in any of the time bins were a decrease in theta power (p=0.04) and an increase in the high gamma power (p=0.02) during the first hour after low-dose DMT infusion (**Figure 6A**). During SWS, theta relative power increased following high-dose DMT (p=0.02) and medium gamma relative power decreased following low-dose DMT (p=0.04) during the first hour post-infusion (**Figure 6B**). DMT infusion at both low and high doses suppressed REM sleep during the first post-infusion hour due to which there was insufficient data for spectral analysis. (**Figure 6C**). The only statistical changes observed in hours 2-4 after DMT infusion were a decrease in REM sleep medium gamma power during hour 2 (p=0.02) and a decrease in REM sleep high gamma power in hour 4 (p=0.04) after low-dose DMT (**Figure 6C**).

**Figure 6.**
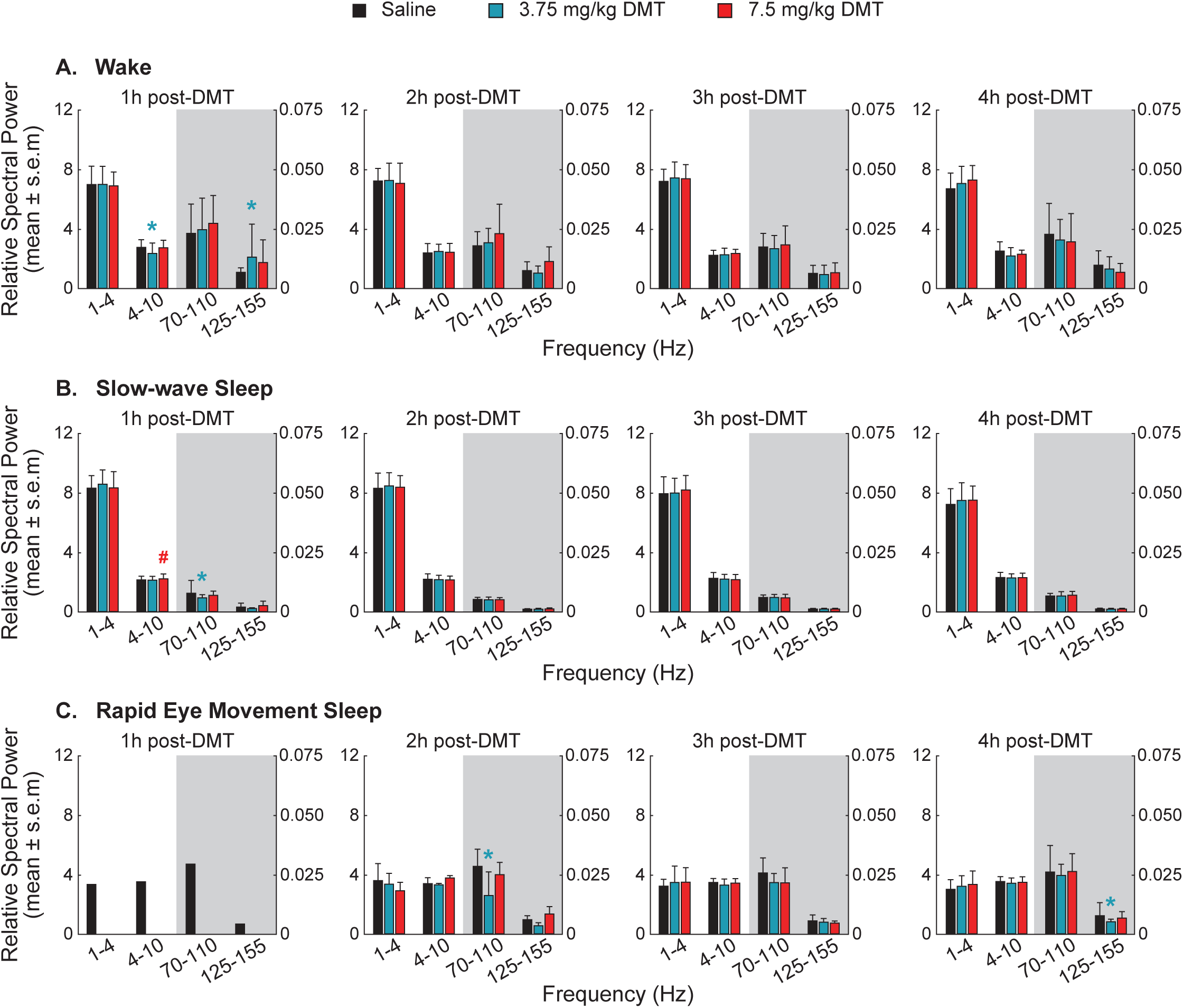
Effect of intravenous administration of N,N-dimethyltryptamine (DMT) on power spectral density across sleep-wake states. Power spectral density analysis of the EEG from first four post-infusion (0.9% saline and DMT) hours. Relative power values were averaged across the delta (1-4 Hz), theta (4-10 Hz), medium gamma (70-110 Hz), and high gamma (125-155 Hz) frequency bands. The effect of intravenous administration of 0.9% saline (black bars), 3.75 mg/kg DMT (teal bars), and 7.5 mg/kg DMT (red bars) on the average global relative power during post-infusion hours 1 through 4 is shown for the wake state (**A.**), slow-wave sleep (**B.**), and rapid eye movement sleep (**C.**) For each plot, the left y-axis corresponds to values for the delta and theta bands whereas the right y-axis corresponds to values for the medium and high gamma bands. The significance symbols denote p<0.05. Asterisk and pound signs indicate statistical significance (p<0.05) for the comparison between low dose DMT (3.75 mg/kg) and saline infusion, and high dose DMT (7.5 mg/kg) and saline infusion, respectively.

### Effect of Intravenous Administration of Psilocybin or DMT on Functional Connectivity Across Sleep-Wake States

As compared to saline infusion, infusion of neither low dose nor high-dose psilocybin produced any statistically significant changes in delta coherence during wake (**Figure 7A**). Infusion of psilocybin at the low dose or the high dose decreased theta coherence and increased medium and high gamma coherence during wake state across the four post-infusion hours (**Figure 7A**). Specifically, low-dose psilocybin decreased theta coherence only in post-infusion hour 1 (p=0.0001) whereas high-dose psilocybin decreased theta coherence across all four hours (hour 1: p=0.0003, hour 2: p=0.003, hour 3: p=0.002, hour 4: p=0.04) (**Figure 7A**). The coherence in the medium gamma frequency band during wake state increased across the four post-infusion hours in both the low-dose (hour 1: p=0.006, hour 2: p=0.0002, hour 3: p=0.004, hour 4: p=0.02) and high-dose (hour 1: p=0.006, hour 2: p<0.0001, hour 3: p=0.0002, hour 4: p<0.0001) psilocybin groups (**Figure 7A**). High gamma coherence during wakefulness increased in all four hours after low dose infusion (hour 1: p<0.0001, hour 2: p<0.0001, hour 3: p=0.0003, hour 4: p<0.0001) and in hour 1 (p<0.0001), hour 2 (p=0.03), and hour 4 (p=0.0007) after high-dose infusion (**Figure 7A**).

**Figure 7.**
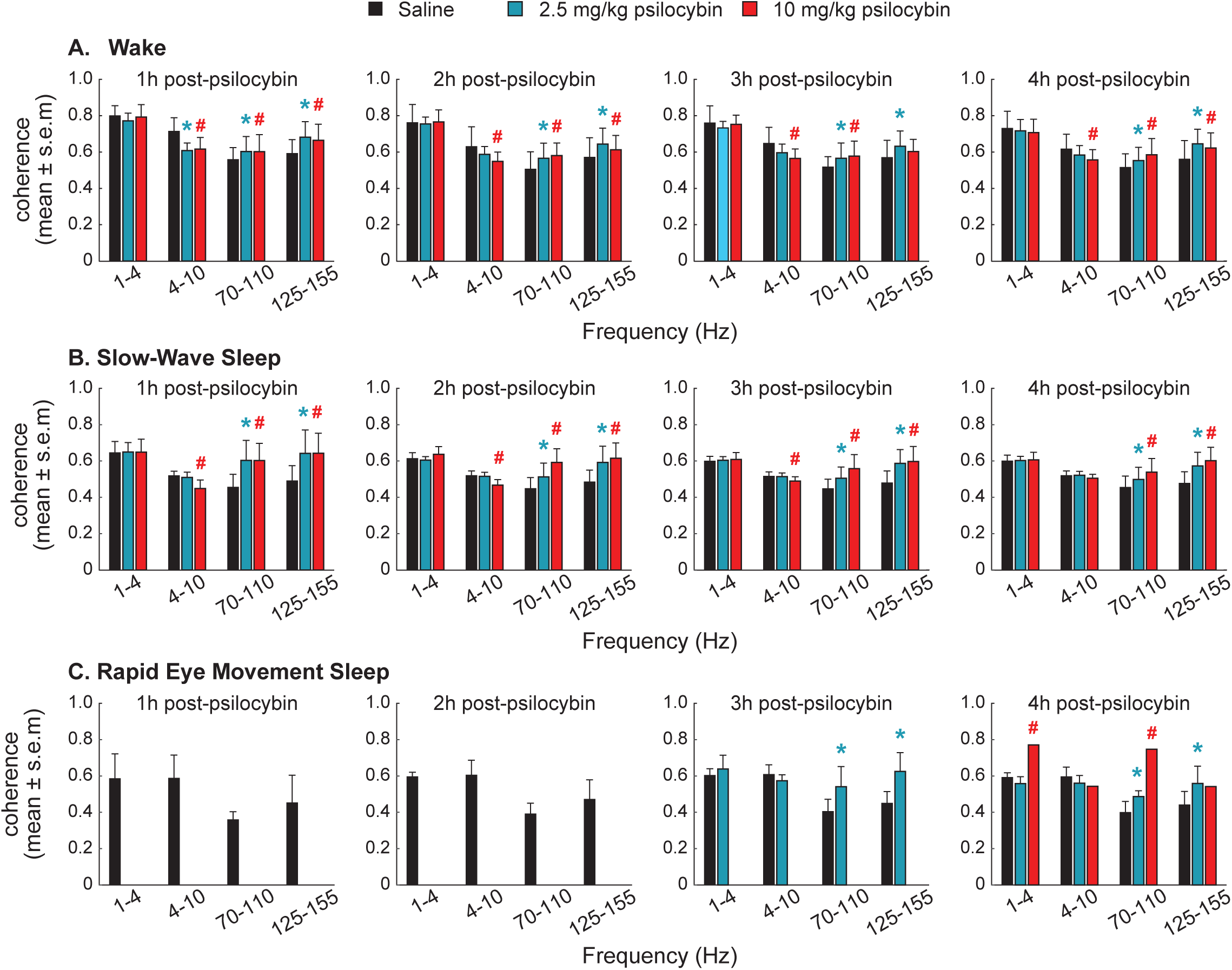
Effect of intravenous administration of psilocybin on corticocortical coherence across sleep-wake states. The EEG data from the first four post-infusion (0.9% saline and psilocybin) hours was analyzed for quantifying the changes in corticocortical coherence, as a surrogate for functional connectivity. Coherence values were averaged across the delta (1-4 Hz), theta (4-10 Hz), medium gamma (70-110 Hz), and high gamma (125-155 Hz) frequency bands. The effect of intravenous administration of 0.9% saline (black bars), 2.5 mg/kg psilocybin (teal bars), and 10 mg/kg psilocybin (red bars) on the global coherence during post-infusion hours 1 through 4 is shown for the wake state (**A.**), slow-wave sleep (**B.**), and rapid eye movement sleep (**C.**) The significance symbols denote p<0.05. Asterisk and pound signs indicate statistical significance (p<0.05) for the comparison between low dose psilocybin (2.5 mg/kg) and saline infusion, and high dose psilocybin (10 mg/kg) and saline infusion, respectively.

During SWS, theta coherence after high-dose psilocybin infusion decreased in hour 1 (p<0.0001), hour 2 (p<0.0001), and hour 3 (p=0.0007) (**Figure 7B**). The coherence in the medium gamma frequency band during SWS increased across all four hours after the infusion of low-(hour 1: p<0.0001, hour 2: p=0.0002, hour 3: p=0.0007, hour 4: p=0.01) or high-dose psilocybin (hour 1: p<0.0001, hour 2: p<0.0001), hour 3: p<0.0001, hour 4: p<0.0001) (**Figure 7B**). During SWS, high gamma coherence increased in all four hours after low-dose (hour 1: p<0.0001, hour 2: p<0.0001, hour 3: p<0.0001, hour 4: p<0.0001) or high-dose (hour 1: p<0.0001, hour 2: p<0.0001, hour 3: p<0.0001, hour 4: p<0.0001) psilocybin infusion (**Figure 7B**).

Due to the suppression of REM sleep after psilocybin infusion, the analyzable REM sleep epochs were found only in hours 3-4 (**Figure 7C**). The medium gamma and high gamma coherence increased during hour 3 (medium gamma: p=0.001; high gamma: p=0.0006) and hour 4 (medium gamma: p=0.02; high gamma: p=0.001) in the low-dose group (**Figure 7C**). In addition, delta (p=0.007) and medium gamma (p<0.0001) coherence were increased during hour 4 in the high-dose psilocybin group (**Figure 7C**).

Infusion of either low-dose or high-dose DMT did not produce any statistically significant changes in delta coherence during wake state (p>0.05) (**Figure 8A**). In contrast, theta coherence during wake state decreased in post-infusion hour 4 for both the low-dose (p=0.02) and high-dose (p=0.02) DMT groups (**Figure 8A**). The medium gamma coherence increased during wake state in the high-dose DMT group during hour 1 (p=0.005), as did high gamma coherence following low-dose (p=0.04) or high-dose (p=0.01) DMT (**Figure 8A**). Infusion of neither low-dose nor high-dose DMT produced any statistically significant effects on coherence during SWS in any of the frequency bands across any of the four time bins except for a decrease in coherence in the delta band after infusion of high-dose DMT during hour 1 (p=0.04) (**Figure 8B**). The only statistically significant changes in coherence during REM sleep during any of the time bins after intravenous DMT were a decrease in theta coherence during hour 2 (p=0.001) in the low-dose DMT group and an increase in theta coherence during hour 4 (p=0.02) in the high-dose DMT group (**Figure 8C**).

**Figure 8.**
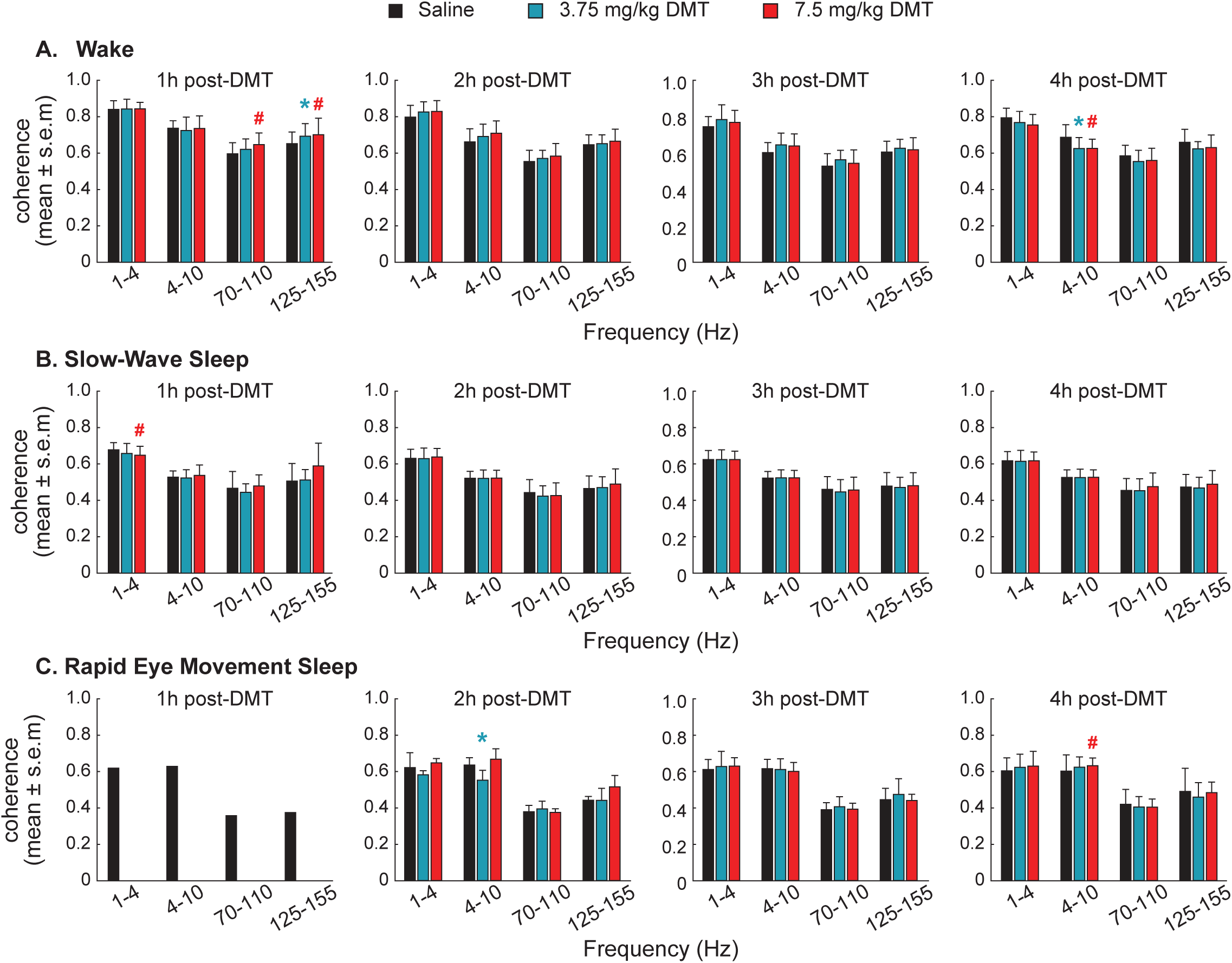
Effect of intravenous administration of N,N-dimethyltryptamine (DMT) on corticocortical coherence across sleep-wake states. The EEG data from the first four post-infusion (0.9% saline and DMT) hours was analyzed for quantifying the changes in corticocortical coherence, as a surrogate for functional connectivity. Coherence values were averaged across the delta (1-4 Hz), theta (4-10 Hz), medium gamma (70-110 Hz), and high gamma (125-155 Hz) frequency bands. The effect of intravenous administration of 0.9% saline (black bars), 3.75 mg/kg DMT (teal bars), and 7.5 mg/kg DMT (red bars) on the global coherence during post-infusion hours 1 through 4 is shown for the wake state (**A.**), slow-wave sleep (**B.**), and rapid eye movement sleep (**C.**) The significance symbols denote p<0.05. Asterisk and pound signs indicate statistical significance (p<0.05) for the comparison between low dose DMT (3.75 mg/kg) and saline infusion, and high dose DMT (7.5 mg/kg) and saline infusion, respectively.

## Discussion

In this study, we demonstrate that intravenous infusion of psilocybin or DMT in rats caused a short-lasting increase in wakefulness, decrease in SWS, and suppression of REM sleep. These data are broadly in agreement with previous studies that showed acute wake-promoting and sleep-suppressing effects of systemic administration of psychedelic 5-HT2A receptor agonists (e.g., LSD, DOI) in rodents.^14,17,19^ Our findings also agree with a more recent mouse study by Thomas and colleagues in which intraperitoneal administration of psilocin, the pharmacologically active metabolite of psilocybin, was shown to cause a short-lasting increase in wakefulness and an increase in the latency to the appearance of REM sleep^18^—an effect that has also been reported in human subjects^13^ and was observed in the current study. These data suggest that any changes in sleep-wake architecture following the administration of psychedelic 5-HT2A receptor agonists are short-lived. Given the common occurrence of sleep disturbances as a co-morbid condition in neuropsychiatric disorders, decreasing the risk of sleep disturbances is vital.^8,9^ Our study, along with the previously published reports in rodents and human subjects, provides key data that alleviates concerns regarding sleep disruptions induced by serotonergic psychedelics, thereby further supporting the therapeutic potential of psychedelic 5-HT2A receptor agonists in treating neuropsychiatric disorders.

The study by Thomas and colleagues also reported an acute increase in spectral power in the delta band of awake animals.^18^ Our results complement this finding, although the enhancement of delta relative power in awake rats was only found in the first hour post-psilocybin infusion and was not observed after DMT administration. In contrast, human studies demonstrate a reduction in delta power after psilocybin administration,^13,23,24^ which has also been reported in select rodent studies.^25^ We also observed a reduction in theta power and theta coherence during wake and SWS in the post-psilocybin period. While the knowledge that such EEG changes persist during SWS is a novel finding in the current study, previous human^23,24,26–29^ and rodent^21,22,25,30,31^ studies have demonstrated a decrease in theta activity during wakefulness following the administration of psilocybin, DMT, and other psychedelic 5-HT2A receptor agonists.

The most salient neurophysiological change after psilocybin and DMT administration was the increase in higher gamma functional connectivity, which was observed across wake, SWS, and REM sleep in the psilocybin group, but was restricted to only during wakefulness in the DMT group. Several rodent studies have reported changes in gamma frequencies during wakefulness after the administration of psychedelic 5-HT2A receptor agonists.^21,22,25,32–34^ While harder to measure in humans due to muscle contamination, similar increases in gamma during wakefulness have been reported after Ayahuasca,^26^ a South American psychoactive beverage with measurable amounts of DMT.^35^ Notably, we report that psilocybin caused changes in theta and high gamma oscillations across sleep-wake states: theta power and coherence decreased during wakefulness and SWS whereas medium and high gamma power and coherence increased across sleep-wake states (i.e., wake, SWS, and REM sleep). Altogether, these data suggest that the psychedelic-induced changes in EEG or neural dynamics can occur independent of the arousal state or the level of arousal.

Although our study assessed two commonly used psychedelics, it is important to determine the effect of other serotonergic psychedelics with longer durations of action (e.g., LSD) as well as atypical psychedelics (e.g., Salvia) on sleep-wake states. Additionally, we administered psilocybin and DMT at the start of the light cycle and therefore cannot comment on the effect of drug administration when performed at different circadian time-points.

In conclusion, as interest in serotonergic psychedelics as therapeutic agents for the treatment of neuropsychiatric disorders grows, and decriminalization efforts expand, it is imperative to rigorously investigate their potential adverse effects.^4–6^ Our data contributes insights into the effects of psilocybin and DMT on sleep-wake states that can inform translational studies aimed at exploring the therapeutic potential of psychedelic 5-HT2A receptor agonists in treating mental health disorders while limiting their impact on sleep.

## Author Contributions

NK, RS, KB, TG, TL performed the experiments, analyzed and interpreted the data; ERH and AN analyzed and interpreted the data; AGH and GV interpreted the data; DP designed the research, analyzed and interpreted the data; all authors contributed to drafting the manuscript.

## Acknowledgements

We thank Chris Andrews, Ph.D. (Consultant, Consulting for Statistics, Computing and Analytics Research core, University of Michigan, 3550 Rackham, 915 East Washington Street, Ann Arbor, MI 48109-1070) for his excellent guidance on statistical analyses. Funding support from the University of Michigan Department of Anesthesiology, NIH R01-GM056398 (AGH), and DMT Quest (DP) is gratefully acknowledged. NK was partially supported by the Rackham Predoctoral Fellowship Award from the Rackham Graduate School at the University of Michigan. ERH was supported by a CTSA Postdoctoral T32 Training Award from the University of Michigan (grant no. T32TR004764).

## Disclosure Statement

The authors declare that this research was conducted in the absence of any conflicts of interest.

## Data Availability Statement

The data used for this manuscript will be made available upon request with a data sharing agreement.

## Notes

### Competing Interest Statement

The authors have declared no competing interest.

